# Evaluating the reproducibility of single-cell gene regulatory network inference algorithms

**DOI:** 10.1101/2020.11.10.375923

**Authors:** Yoonjee Kang, Denis Thieffry, Laura Cantini

## Abstract

Networks are powerful tools to represent and investigate biological systems. The development of algorithms inferring regulatory interactions from functional genomics data has been an active area of research. With the advent of single-cell RNA-seq data (scRNA-seq), numerous methods specifically designed to take advantage of single-cell datasets have been proposed. However, published benchmarks on single-cell network inference are mostly based on simulated data. Once applied to real data, these benchmarks take into account only a small set of genes and only compare the inferred networks with an imposed ground-truth.

Here, we benchmark four single-cell network inference methods based on their reproducibility, i.e. their ability to infer similar networks when applied to two independent datasets for the same biological condition. We tested each of these methods on real data from three biological conditions: human retina, T-cells in colorectal cancer, and human hematopoiesis.

GENIE3 results to be the most reproducible algorithm, independently from the single-cell sequencing platform, the cell type annotation system, the number of cells constituting the dataset, or the thresholding applied to the links of the inferred networks. In order to ensure the reproducibility and ease extensions of this benchmark study, we implemented all the analyses in scNET, a Jupyter notebook available at https://github.com/ComputationalSystemsBiology/scNET.

## 1 Introduction

Biological systems are inherently complex, in particular because of the emergent phenotypic properties arising from the interaction of their numerous molecular components. Characterizing genotype to phenotype connections and deregulations toward disease thus requires to identify the biological macromolecules involved (e.g. genes, mRNAs, proteins), but also how these interact in a huge diversity of cellular pathways and networks (Barabási and Oltvai, 2004).

In the post-genomic era, biological networks have been extensively exploited to investigate such complex interactions among biological macromolecules (Barabási et al., 2011; Sonawane et al., 2019; Silverman et al., 2020). Network-based studies brought crucial insights into cell functioning and diseases (Basso et al., 2005; Margolin et al., 2006; Ideker and Sharan, 2008). A network is a graph-based representation of a biological system, where the nodes represent objects of interest (e.g. genes, mRNAs, proteins), while the edges represent relations between these objects (e.g. gene co-expression, or binding between two proteins). Different approaches can be used to reconstruct biological networks. Here, we focus on data-driven methods, which infer networks from gene expression data with the help of reverse engineering techniques (Sonawane et al., 2019).

Network inference algorithms were first proposed to extract information from bulk gene expression data, and their development has been an active area of research for more than 20 years (Barabási et al., 2011; Verny et al., 2017; Sonawane et al., 2019; Silverman et al., 2020). With the advent of single-cell RNA sequencing (scRNA-seq), we started to gather transcriptomic data from individual cells, enabling proper studies of their heterogeneity. However, the analysis of scRNA-seq data comes with a variety of computational challenges (e.g. small number of sequencing reads, systematic noise due to the stochasticity of gene expression at single-cell level, dropouts) that distinguish this data type from its bulk counterpart. For this reason, network inference methods originally developed for bulk gene expression data may not be suitable for data generated from single cells. The development of network inference algorithms has thus recently undergone a strong shift towards the design of methods targeting single-cell data (Fiers et al., 2018).

Two benchmarks of single-cell network inference methods have been published (Chen and Mar, 2018; Pratapa et al., 2020). Both works evaluate network inference algorithms by comparing the inferred network with a ground-truth. These works are also mostly focused on simulated data and they apply a strong filtering on genes (leaving only 100-1,000 genes for network inference). Chen et al. (Chen and Mar, 2018) considered five methods targeting bulk data and three methods specifically designed for single-cell data. More recently, Paratapa et al. (Pratapa et al., 2020) focused on twelve methods designed for single-cell data. Both benchmarks concluded that the overall performances of all methods were quite disappointing, and that network inference remains a challenging problem.

Here, we evaluate network inference algorithms based on their reproducibility, i.e. their ability to infer similar networks once applied to two independent datasets for the same biological condition (e.g. two independent scRNA-seq datasets of colorectal cancer). The rationale behind this comparison is that, if the two independent datasets are profiled from the same biological condition (e.g. colorectal cancer) involving the same cell types, we can expect that the regulatory programs underlying them should strongly overlap. As a consequence, a good network inference algorithm should infer highly overlapping networks when applied to single-cell datasets profiled from the same biological condition. Starting from the work of Paratapa et al., we selected the four algorithms that do not require an ordering of the cells according to pseudo-time and we tested the reproducibility of the inferred networks in three biological systems: human retina, T-cells in colorectal cancer and human hematopoiesis. Differently from previous benchmarks, we only applied a soft filtering on genes, thus testing the algorithms based on their performances to infer networks involving from 6000 to 12000 nodes/genes.

From our benchmark, GENIE3 emerges as the most reproducible network inference algorithm. Interestingly this performance is not influenced by the single-cell sequencing platform, the cell type annotation system, the number of cells constituting the single-cell dataset, or the thresholding applied to the links of the inferred networks. In order to ensure the reproducibility and ease extensions of this benchmark study, we implemented all the analyses in a Jupyter notebook, called scNET and available at https://github.com/ComputationalSystemsBiology/scNET.

## 2 Materials and Methods

### 2.1 Single-cell network inference algorithms benchmarked

Starting from the exhaustive collection of single-cell network inference algorithms presented in (Pratapa et al., 2020), two main categories of methods can be distinguished. Some methods interpret scRNA-Seq as time-course expression data, where the pseudo-time corresponds to the time information. These methods are frequently based on Ordinary Differential Equations (ODEs) and are relevant for biological systems undergoing dynamic transcriptional changes (e.g. scRNA-Seq performed on differentiating cells) (Matsumoto et al., 2017). In contrast, other methods do not use pseudo-time information to infer networks. These methods generally use statistical measures (e.g. correlation, mutual information) to infer regulatory connections and are thus better suited for transcriptomic data not affected by strong dynamical processes (e.g. retina cells in normal state).

Testing reproducibility strictly requires the availability of two independent scRNA-seq datasets reflecting the same biological condition and presenting as few as possible technical variations. Indeed, the presence of technical variations due to the sequencing or experimental procedures could drastically impact the conclusions of our work. In this respect, finding independent scRNA-seq datasets reflecting dynamic transcriptional changes, generated with the same experimental procedure, is really challenging. We thus decided to focus our benchmark study on network inference methods that do not use the pseudo-time information. Four single-cell network inference methods are thus considered in this evaluation: GENIE3 (Huynh-Thu et al., 2010), GRNBoost2 (Moerman et al., 2019), PIDC (Chan et al., 2017) and PPCOR (Kim, 2015). Of note, the first three algorithms are also the best performing in the benchmark of Pratapa et al.

<u>GEne Network Inference with Ensemble of Trees (GENIE3)</u> (Huynh-Thu et al., 2010) is a tree-based network inference method. For each gene G_1_ in the expression dataset, GENIE3 solves a regression problem, determining the subset of genes whose expression is the most predictive of the expression of G_1_. This method was the best performing algorithm in the DREAM4 In Silico Multifactorial challenge (Greenfield et al., 2010). GENIE3 requires in input the scRNA-seq expression matrix and a list of Transcription Factors (TFs). In our tests the list of human TFs provided in input corresponds to the intersection between the expressed genes and those annotated as encoding TFs by (Chawla et al., 2013). The output of GENIE3 is a weighted network linking TFs with predicted target genes. The weight associated with each link corresponds to its Importance Measure (IM), which represents the weight that the Transcription Factor has in the prediction of the level of expression of the target gene. No post processing threshold has been applied to the inferred links.

<u>GRNBoost2</u> (Moerman et al., 2019) has been developed as a faster alternative to GENIE3. It is thus based on a regression model, using a stochastic gradient boosting machine regression. The inputs and outputs of GRNBoost2 are the same as for GENIE3, and no post processing threshold has been applied to the inferred links. Both GRNBoost2 and GENIE3 are part of the SCENIC workflow (Aibar et al., 2017).

<u>PPCOR</u> (Kim, 2015) infers the presence of a regulatory interaction between two genes by computing the correlation of their expression patterns. To control for possible indirect effects, partial correlation is used instead of a simple correlation, where partial correlation is a measure of the relationship between two variables while controlling for the effect of other variables. The only input of PPCOR is the expression matrix. The output of PPCOR is a weighted network, where all links are weighted based on the partial correlation between the expression values of the linked nodes/genes. The network produced by PPCOR is complete, i.e. all nodes are connected with all. We thus had to filter the links of the inferred network based on the significance of the correlation values associated to the links (P-value threshold 0.05).

<u>Partial Information Decomposition and Context (PIDC)</u> (Chan et al., 2017) is based on concepts from information theory and uses partial information decomposition (PID) to identify potential regulatory relationships between genes. The only input of PPCOR is the expression matrix and its output is a weighted gene-gene network.

### 2.2 Data acquisition and preprocessing

Fourteen public scRNA-seq datasets have been used for this benchmark: Menon and Lukowski obtained by profiling huma retina cells; Zhang and Li profiling T-cells in colorectal cancer (CRC); Hay and Setty profiling human hematopoiesis cells. See Table 1 for a complete description of these datasets. The hematopoiesis datasets were split according to their cell type of origin. Only those cell types reported in both studies by Hay et al. and Setty et al. were considered. We thus obtained a total of 10 scRNA-seq datasets in hematopoiesis spanning five cell types: HSC, CLP, Monocyte, Erythroblast and Dendritic Cell.

**Table 1.**
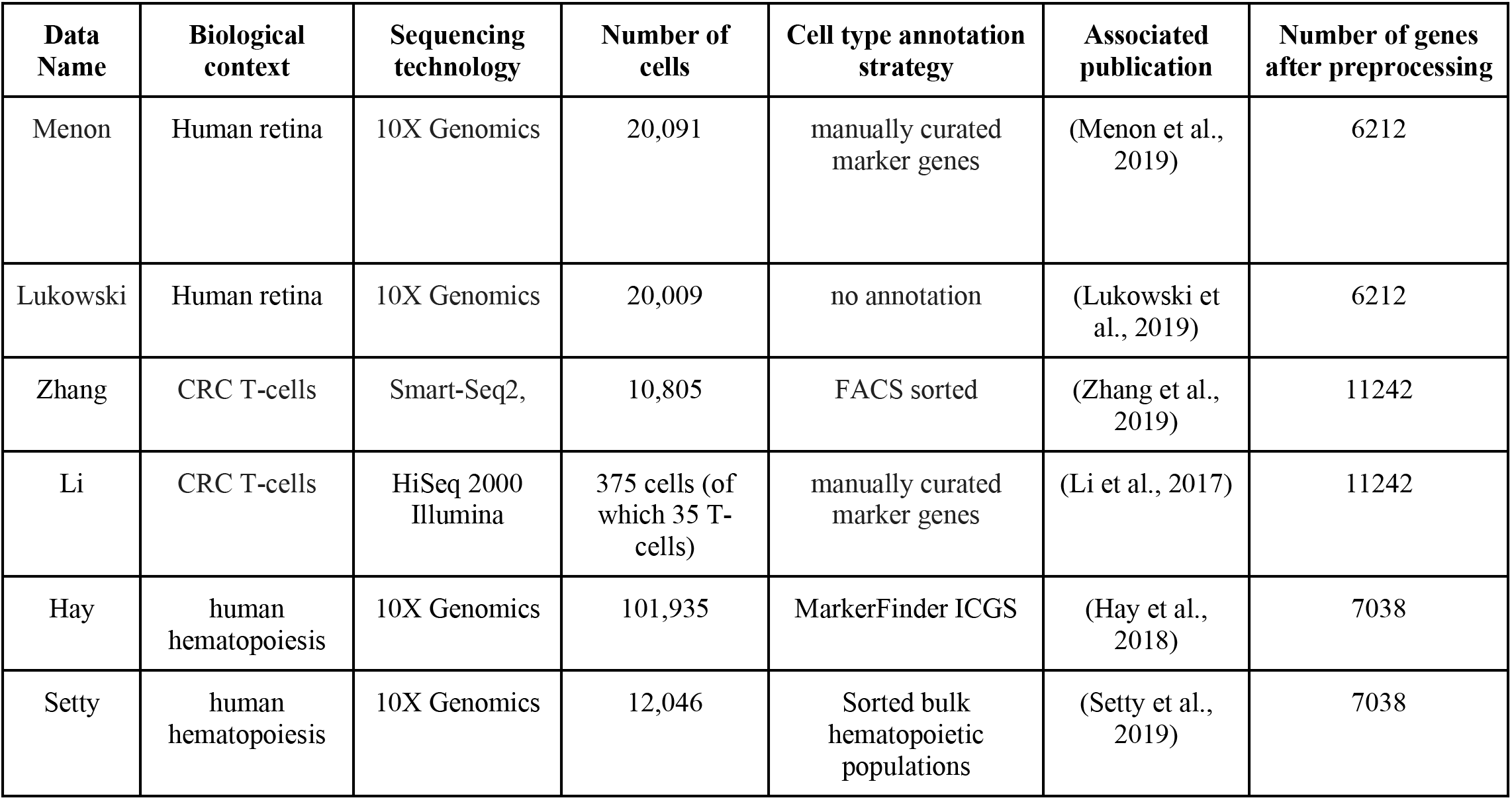
Datasets employed in this benchmark.

After downloading the data, we filtered the genes based on their total count number (< 3 *0.01*number of cells), as well as on the number of cells in which they are detected (>0.01*number of cells), as described in (Aibar et al., 2017). The gene filtering is performed on each dataset independently. Then, for each biological condition (CRC T-cells, retina, hematopoiesis), only the genes retained for both datasets were selected for network inference. The number of genes retained after filtering are reported in the last column of Table 1. Finally, the data were log2-normalised before applying the different network inference algorithms.

### 2.3 Indexes employed to measure the reproducibility of the network inference algorithms

Percentage of intersection (perINT) and Weighted Jaccard Similarity (WJS) have been employed here to test the reproducibility of the network inference algorithms. The percentage of intersection is used to detect the presence of links shared between two compared networks, while WJS takes into account the similarity of the weights associated with the links shared between the compared networks.

Given two networks N1 and N2 inferred respectively from scRNAseq datasets D1 and D2, and indicating as |*N*| the number of links in the network N, the percentage of intersection (perINT) is computed as:

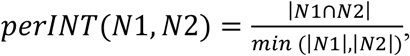

while the Weighted Jaccard Similarity (WJS) (Tantardini et al., 2019), is defined as

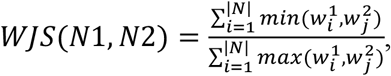

where *w*^1^, *w*^2^ are the vectors of weights associated with the links in common between N1 and N2.

In addition, to compare the inferred links to a ground-truth, we also considered a RcisTarget score derived from the application of the RcisTarget tool (Aerts et al., 2010; Aibar et al., 2017). Given a network of TF-gene interaction, RcisTarget predicts candidate target genes of a TF by looking at the DNA motifs that are significantly over-represented in the surroundings of the Transcription Start Site (TSS) of all the genes that are linked to the TF. We here consider the links validated by RcisTarget as ground-truth and we compare them with the inferred networks, by computing:

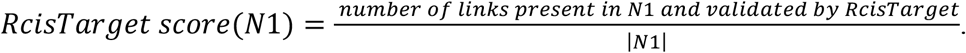

In the case of the methods inferring links between all genes, a selection of links connecting TFs with possible target genes is performed before computing the RcisTaget score.

### 2.4 Testing if the number of links in the networks affects our reproducibility score

The number of links inferred by the network inference algorithm can affect our reproducibility tests. For example, in the extreme case of a method inferring complete networks, the perINT score would be 100%. To test whether our results were affected by the number of links inferred by the different methods, we constructed a null model. Starting from the two networks inferred in a given biological condition (e.g. human retina), we randomly reshuffled the links of the two networks independently and tested the reproducibility scores. The reshuffling of the links in GENIE3 and GRNBoost2 was realized taking into account the different roles played by TFs and the other genes in the network. After repeating this procedure 10,000 times, we could verify the positioning of the real reproducibility scores with respect to the distribution obtained with the null model, and thereby assign p-values to the scores.

## 3 Results

Starting from the work (Pratapa et al., 2020) we selected the four single-cell network inference algorithms that do not require an ordering of the cells according to pseudo-time (GENIE3, GRNBoost2, PPCOR and PIDC, see Materials and Methods) and we evaluated them based on their reproducibility, i.e. their ability to infer similar networks once applied to two independent datasets from the same biological condition (e.g. two independent scRNA-seq datasets of colorectal cancer). The reproducibility is measured based on Percentage of intersection (perINT) and Weighted Jaccard Similarity (WJS) (see Materials and Methods). In addition, we computed the intersection with a ground-truth, based on the RcisTarget score (see Materials and Methods). The evaluation is repeated across three biological conditions: human retina, T-cells in colorectal cancer and human hematopoiesis, for a total of fourteen independent scRNAseq datasets. See Figure 1 for an overview of the benchmark workflow.

**Figure 1.**
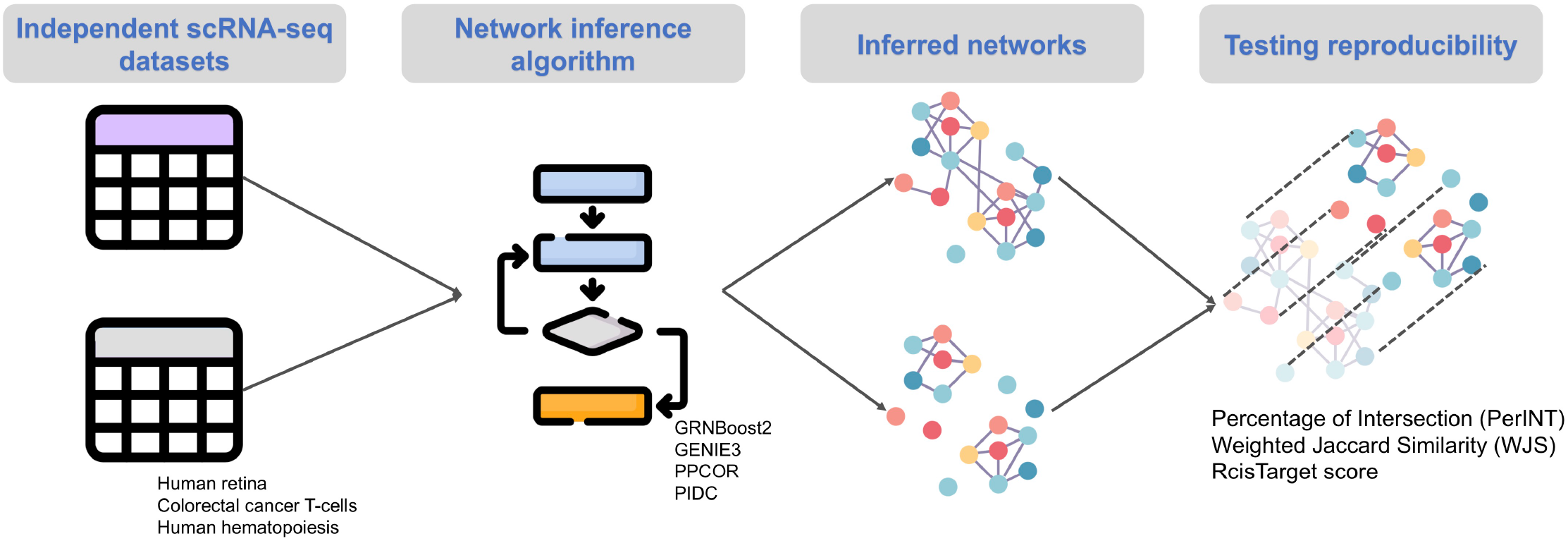
Summary of the workflow followed in this benchmark.

While in previous benchmarks (Chen and Mar, 2018; Pratapa et al., 2020) a low number of highly variable genes had been taken into account (100-1000 genes), we here tested the ability of the algorithms to infer networks involving all expressed genes (see Materials and Methods for details on the procedure used to filter genes). Indeed, filtering only the top 100-1,000 varying genes is a strong limitation. Restricting the nodes of the inferred network to a low number of genes is reasonable when a manually curated list of relevant genes is available (for example marker genes identified by wet-lab experiments). However, when such a list is not available, working only with the top 100-1000 varying genes may overlook genes and interactions playing a key role in the regulatory programs of the biological system. We thus tested the various network inference algorithms once applied to scRNAseq datasets containing 6,000-11,000 genes.

In our test cases, PIDC failed to reconstruct the networks for two main reasons: (i) the algorithms was slow, especially in the discretization step required to infer the network, and (ii) the use of multivariate information measures impose to have a number of genes much lower than the number of cells, thus requiring to drastically filter out the starting set of genes. Overall, PIDC thus resulted to be more adequate to infer small networks (100-1,000 nodes/genes), which are not the focus of this work.

### 3.1 Reproducibility in human retina

We applied GENIE3, GRNBoost2 and PPCOR to two independent scRNA-seq datasets of human retina, reported in (Menon et al., 2019) and in (Lukowsk et al., 2019) (see Materials and Methods). After filtering, the two datasets span 6,212 common genes across a comparable number of cells: 20,091 in Menon versus 20,009 in Lukowski.

We thus inferred a total of six networks. Of note, similar network sizes were obtained across the three network inference algorithms and across datasets, encompassing approximately one million links each (see Supplementary Table 1 for details). We then evaluated the reproducibility of each algorithm by computing the Percentage of intersection (perINT) and the Weighted Jaccard Similarity (WJS) between the networks inferred independently from the two datasets The percentage of intersection is intended to test the amount of common links between the two networks, while the WJS takes also into account the similarity of the weights associated with the common links.

As shown in Figure 2A, GENIE3 is the algorithm showing the highest reproducibility according to both indexes, with a perINT reaching 100% and a WJS at 0.67. Our null model confirms that these results are not affected by the number of inferred links (see Materials and Methods for further details and Supplementary Table 2 for the corresponding P-values). At the same time, in agreement with the results of the previous benchmarks, the intersection with the ground true considered remains rather low, with RcisTarget scores ranging within 0-1.9%.

**Figure 2.**
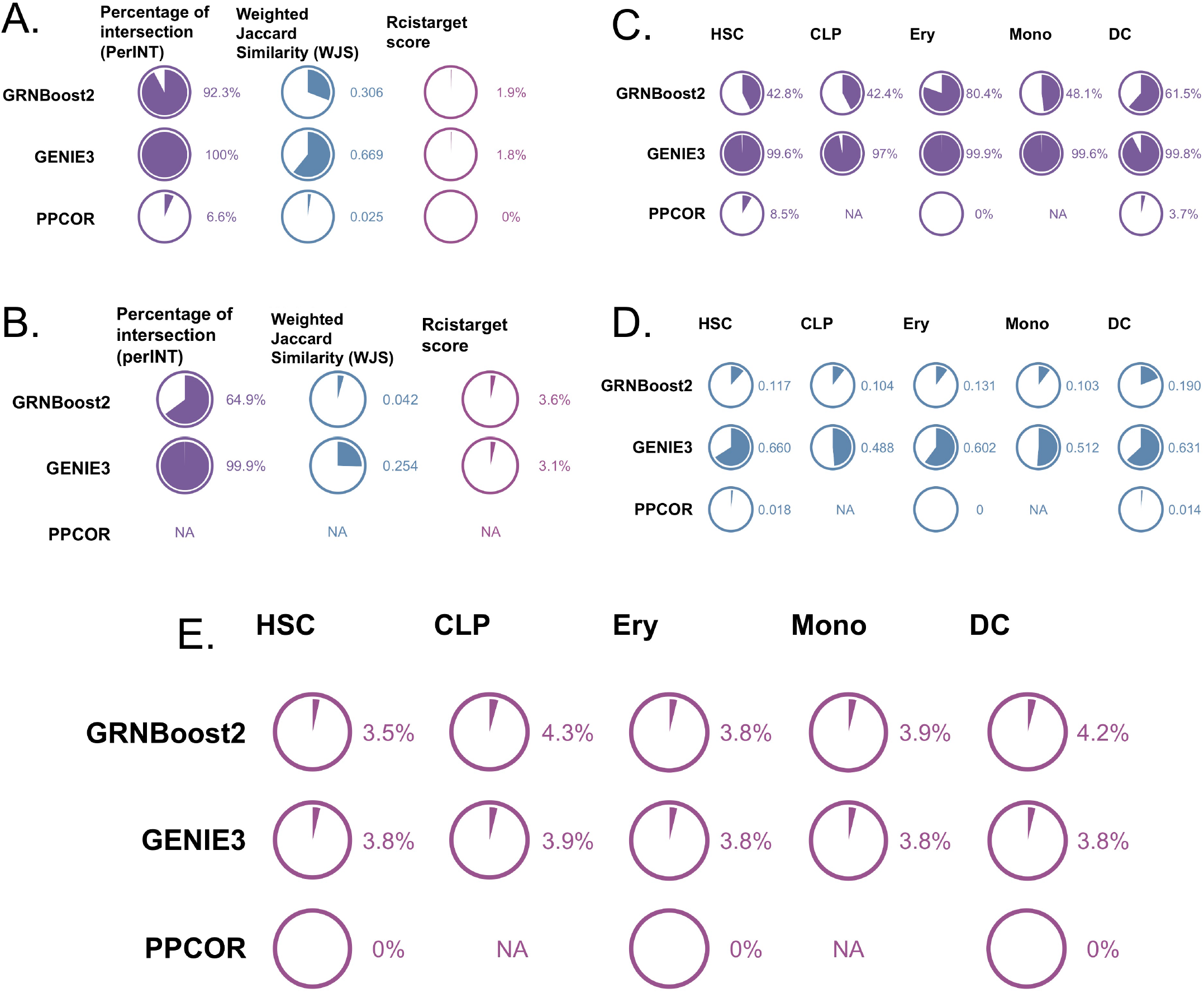
Reproducibility performances of the various network inference algorithms across the three biological contexts: human retina, colorectal cancer T-cells and human hematopoiesis. A and B report summarise the Percentage of intersection (perINT), Weighted Jaccard Similarity (WJS) and RcisTarget score obtained by the benchmarked algorithms (GRNBoost2, GENIE3 and PPCOR) in human retina and colorectal cancer T-cells respectively. C-E summarize the performances of the same algorithms in hematopoiesis, with perINT (in C), WJS (in D) and RcisTarget score (in E).

### 3.2 Reproducibility in colorectal cancer (CRC) T-cells

We further tested the performances of GENIE3, GRNBoost2 and PPCOR in colorectal cancer (CRC) T-cells. The two datasets used in this case are taken from (Zhang et al., 2019) and (Li et al., 2017) (see Materials and Methods), restricting the last dataset to only T-cells (see Materials and Methods). After filtering, we obtained datasets composed of 11,242 common genes and a widely varying number of cells: 10,805 for Zhang and 35 for Li.

Applying GENIE3, GRNBoost2 and PPCOR independently to the two datasets, we observe a high variability in the number of inferred links, which tend to be much lower in Li et al. compared to Zhang et al., presumably due to the high difference in the number of cells profiled in the two datasets (see Supplementary Table 2 for details). At the same time, variations across algorithms could be also observed, with GENIE3 inferring the highest number of links (three million and six million in Li and Zhang, respectively). Of note, PPCOR has been excluded from this comparison, as it produced partial correlation values outside the range [−1;1] for the Li et al. dataset.

After computation of the perINT and WJS (Figure 2B), GENIE3 emerged as the best performing method, with a perINT of 99.9% and a WJS of 0.25. Our null model confirms that these results are not affected by the higher number of links inferred by GENIE3 (see Materials and Methods for further details and Supplementary Table 2 for the corresponding P-values). Also, in this case, the RcisTarget score reflecting the intersection with a ground-truth is quite low (3.1-3.6%). Of note, despite the low number of cells reported by Li, the RcisTarget score obtained in this dataset is comparable with those obtained in networks inferred from much larger datasets.

### 3.3 Reproducibility in human Hematopoiesis

Human hematopoiesis has been used as the third biological context for the comparison of GENIE3, GRNBoost2 and PPCOR. The hematopoiesis datasets were split according to the different cell types profiled: HSC, CLP, Monocyte, Erythroblast and Dendritic Cell, obtaining a total of 10 scRNA-seq datasets. Networks were thus inferred on each cell type independently with GENIE3, GRNBoost2 and PPCOR, resulting in a total of 30 networks. Also, in this case, GENIE3 led to the highest number of links (approximately 2 million in all cell types), while GRNBoost2 and PPCOR led to numbers of links varying from 700 thousands to one million (see Supplementary Table 1). As for CRC T-cells, PPCOR produced networks composed of links with partial correlation higher than 1 and/or lower than −1 for some CLPs, and Monocytes. For this reason, we did not consider PPCOR in the reproducibility evaluation for these cell types.

The reproducibility was then tested for each cell type using the perINT and WJS indexes (Figure 2C-D). Here also, GENIE3 displayed the best performances with percentages of intersection reaching 97-100% and WJS at 0.5-0.66. Our null model confirms that these results are not affected by the higher number of links inferred by GENIE3 (see Materials and Methods for further details and Supplementary Table 2 for the obtained P-values). Consistently with previous observations, the RcisTarget scores remains low (3.5-4.3%) for all cell types and all methods (Figure 2E).

### 3.4 Stability with respect to link thresholding in the inferred networks

All the networks inferred by GENIE3, GRNBoost2 and PPCOR could be thresholded based on the distribution of the weights associated with their links. In the results presented above, the networks inferred with GENIE3 and GRNBoost2 did not undergo any filtering, given that these tools already perform a selection on the links. In contrast, the networks obtained with PPCOR are complete (i.e. everything is connected with everything), calling for a filtering of the links, which was done based on the significance of the correlation values (see Materials and Methods).

To test if more stringent filtering could alter our conclusions regarding the reproducibility of the benchmarked methods, we filtered the links of the inferred networks based on the distribution of the weights of these links. For all network inference methods, we imposed three thresholds on the weight distribution of the links, retaining the 40th, 80th and 90th percentiles. After thresholding, the intersection between the networks inferred on independent datasets from the same biological condition were evaluated, using the percINT and WJS as above.

As shown in Figure 3, the performances of all network inference methods tend to decrease when the threshold is increased, suggesting that the weight of the links is not a good proxy for their reproducibility. Overall, GENIE3 remains the best performing method independently on the threshold employed.

**Figure 3.**
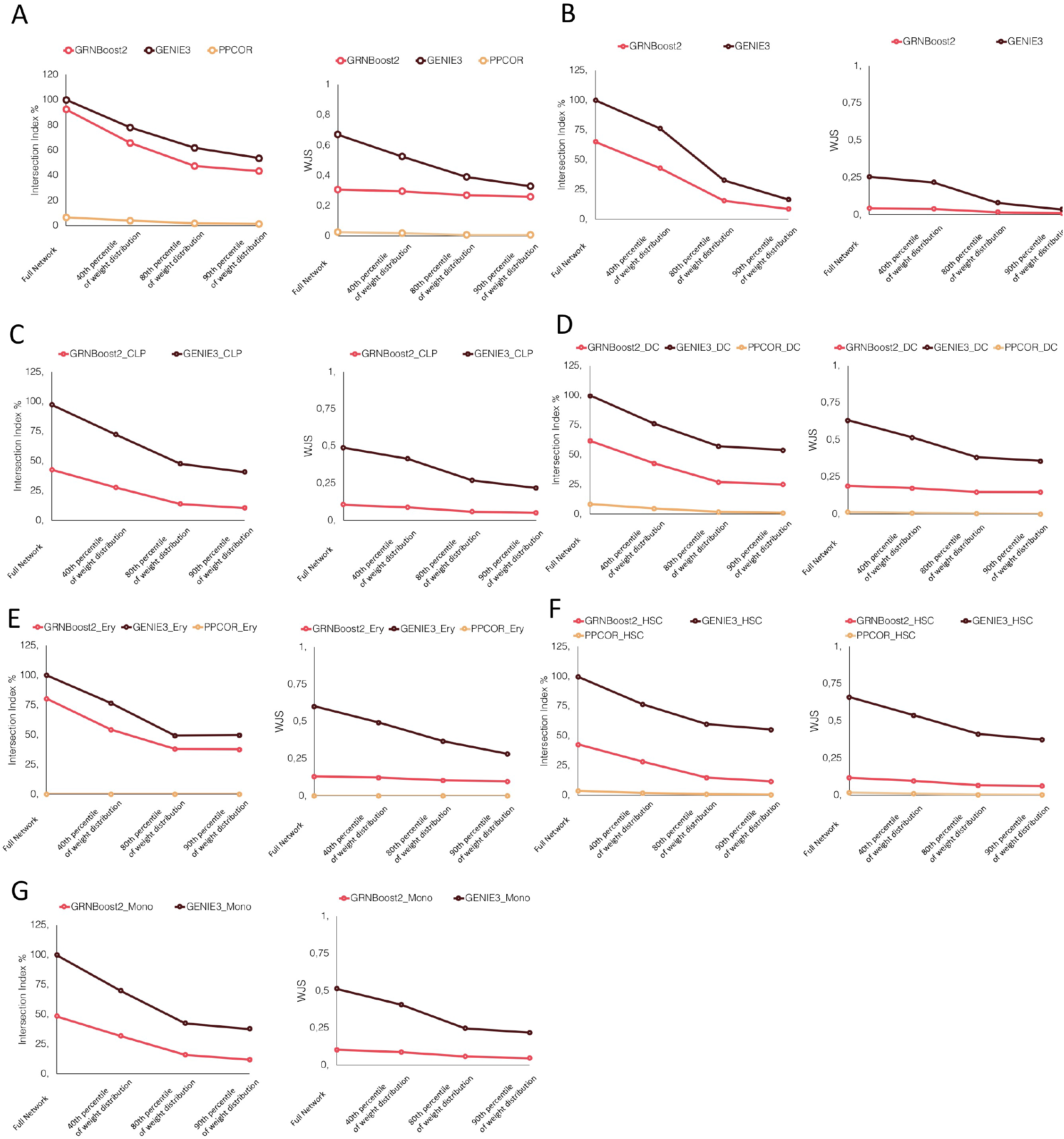
perINT and WJS according to different network thresholding. The perINT and WJS are here reported for varying thresholds on the weight distribution of the links of the inferred networks. THe results are reported for all the tested datasets (A) retina, (B) CRC T-cells, (C) CLPs, (D) Dendritic cells, (E) Erythrocytes, (F) HSCs, (G) Monocytes.

### 3.5 Stability with respect to technical variations in the input data: number of profiled cells, sequencing platform and cell type annotation

In the experiments performed above, we tested the reproducibility of the network inference algorithms by using two independent datasets for each biological condition (e.g. human retina). A limitation of this approach comes from the technical differences between the protocols followed to generate these datasets: different sequencing platforms, different procedures used for the annotation of the cell types, and different number of cells. All these technical differences could impact our results.

To evaluate the stability of the results against technical variations, we used the largest dataset, from (Menon et al., 2019), encompassing 20,091 cells. We splitted this dataset into two subsets, keeping the proportions of the various cell types constant. We then applied the three network inference algorithms independently to the two subsets and we evaluated the reproducibility of the algorithms using perINT and WJS, as in the previous tests. To further assess the effect of the number of cells on network inference, we split the same scRNAseq dataset generated by Menon et al. three times to obtain couples of datasets encompassing decreasing number of cells: 100,000,1,000 and 100. Note that for all these comparisons, the sequencing platform and/or the method/technique used to annotate the cells are identical for all subsets

PPCOR inferred networks for 100,000 and 1,000 cells, but failed at 100 cells by displaying correlation values outside the range [−1;1] (see Supplementary Table 3). In addition, as shown in Figure 4, GENIE 3 emerged again as the best performing method in all cases. Of note, when varying the number of cells in the input data, the percentage of intersection and the number of links barely vary (see Figure 4 and Supplementary Table 3), while the WJS decreases more drastically (from 0.8 to 0.3 for GENIE3).

**Figure 4.**
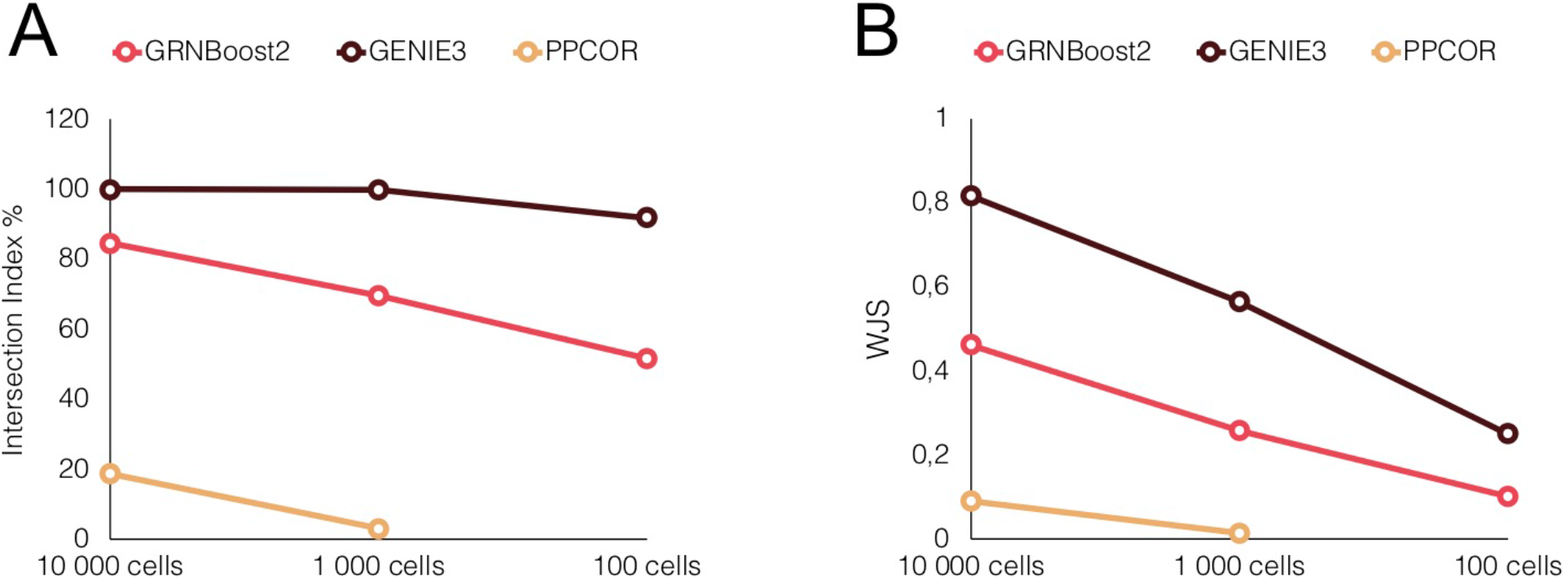
Stability of the network inference performances with respect to technical variations in the input data. Reproducibility scores of GRENBoost2 (red), GENIE3 (black) and PPCOR (yellow) across different splittings of the Menon, M. et al. retina dataset. A and B correspond to the percentage of intersection (perINT) and Weighted Jaccard Similarity (WJS), respectively.

### 3.6 The scNET Jupyter notebook

To foster the reproducibility of all the results and figures presented in this study, we implemented the corresponding code in a Jupyter notebook available at https://github.com/ComputationalSystemsBiology/scNET together with the associated Conda environment containing all the required libraries installed. Importantly, scNET can be used to benchmark new network inference algorithms based on their reproducibility, or further test GENIE3, PPCOR and GRNBoost2 on user-provided datasets.

## 4 Discussion

Starting from the benchmark of Paratapa et al., we evaluated the network inference algorithms from a complementary perspective by assessing their reproducibility. We were thus interested to test if the algorithms would infer the same network once applied to pairs of independent datasets from the same biological condition (e.g. T-cells in colorectal cancer). Our benchmark focused on real patient-derived data spanning three biological contexts: human retina, T-cells in CRC, and human hematopoiesis cells. We thus span highly different biological contexts, going from cancer tissue, to isolated healthy immune cells, and to a mixture of normal retina cells combined in a single dataset. Importantly, we aimed at inferring networks involving a much higher number of genes compared to previous works.

In agreement with previous benchmarks, all network inference algorithms generated networks having low intersections with ground-truth. Of note the ground-truth considered here, based on RcisTarget, is different and complementary to those used in previous benchmarks. This disappointing result might arise for different reasons, potentially adding up. Limitations can be present in the input data, as scRNAseq may not provide sufficient resolution for reliable network inference. Turning to the inference algorithm, limitations may arise from underlying statistical assumptions. Finally, the ground-truth network considered here and in previous benchmarks may not be sufficiently comprehensive.

GENIE3 consistently generated the most reproducible results across all the three biological contexts considered. Furthermore, its performances proved to be stable with respect to the single-cell sequencing platform, the cell type annotation system, the number of cells considered as well as with respect to the thresholding applied to the links of the inferred networks. PPCOR provided values outside the normal range of correlation values ([−1,1]) for datasets having less than 1000 cells. Such inconsistencies are likely due to numerical problems arising when the input dataset encompasses many more genes than cells.

The main limitation of this benchmark is the number of considered network inference algorithms. Future extensions of this study could include pseudotime-based network inference methods, once adequate datasets will become available. To date, available independent datasets relevant for pseudotime-based network inference algorithms (e.g. cells profiled during development stimulation) present too many experimental variations to be employed for a reliable evaluation of reproducibility. Of note, such extensions will be greatly facilitated by taking advantage of the Jupyter notebook (scNET) provided as supplementary material.

## Conflict of Interest

The authors declare that the research was conducted in the absence of any commercial or financial relationships that could be construed as a potential conflict of interest.

## Author Contributions

LC designed the analysis. YK performed the analysis. LC and DT co-supervised the study. All authors contributed to the manuscript and approved the submitted version.

## Acknowledgments

We thank the bioinformatics platform of IBENS for the computational/infrastructural support. We thank Michael Mason, Anaïs Baudot and Sabine Tejpar for the scientific feedbacks on the work.

## Data Availability Statement

The datasets for this study can be accessed from their associated publications (see Table1). All the analyses are reproducible using the scNET Jupyter notebook available at https://github.com/ComputationalSystemsBiology/scNET.

## Supplementary Material

### 1 Supplementary Tables

**Supplementary Table 1.** Number of links in the various inferred networks.

|  |  |  |
| --- | --- | --- |
| Data Name | Algorithm | Number of links |
| Menon et al. | PPCOR | 598539 |
|  | GENIE3 | 1552750 |
|  | GRNBoost2 | 1421357 |
| Lukowski et al. | PPCOR | 1184848 |
|  | GENIE3 | 1552750 |
|  | GRNBoost2 | 1355892 |
| Zhang et al. | PPCOR | 1237822 |
|  | GENIE3 | 5833037 |
|  | GRNBoost2 | 3006644 |
| Li et al. | PPCOR | NA |
|  | GENIE3 | 2950874 |
|  | GRNBoost2 | 765846 |
| Hay et al. HSC | PPCOR | 571935 |
|  | GENIE3 | 2448676 |
|  | GRNBoost2 | 801816 |
| Hay et al. CLP | PPCOR | NA |
|  | GENIE3 | 2321809 |
|  | GRNBoost2 | 764244 |
| Hay et al. Monocytes | PPCOR | NA |
|  | GENIE3 | 2418779 |
|  | GRNBoost2 | 799381 |
| Hay et al. Erythroblast | PPCOR | 761300 |
|  | GENIE3 | 2461623 |
|  | GRNBoost2 | 1787691 |
| Hay et al. Dendritic Cell | PPCOR | 1703169 |
|  | GENIE3 | 2453534 |
|  | GRNBoost2 | 1184762 |
| Setty et al. HSC | PPCOR | 566853 |
|  | GENIE3 | 2447457 |
|  | GRNBoost2 | 726544 |
| Setty et al. CLP | PPCOR | NA |
|  | GENIE3 | 2332534 |
|  | GRNBoost2 | 607112 |
| Setty et al. Monocytes | PPCOR | 514936 |
|  | GENIE3 | 2452913 |
|  | GRNBoost2 | 962318 |
| Setty et al. Erythroblast | PPCOR | 249941 |
|  | GENIE3 | 2448696 |
|  | GRNBoost2 | 1143651 |
| Setty et al. Dendritic Cell | PPCOR | 360772 |
|  | GENIE3 | 2457673 |
|  | GRNBoost2 | 1265417 |

**Supplementary Table 2.** P-values null model. The perINT index of our experiments are here compared in respect to the distribution of perINT indexes obtained over 1000 random reshufflings of the networks. The value ≤0.001 correspond to a zero over 1000 runs, which indicates a P-value lower than 0.001.

| Data Name | Algorithm | P-value null model |
| --- | --- | --- |
| Retina Manon et al. and Lukowski et al. | PPCOR | 0.176 |
| | GENIE3 | $\leq 0.001$ |
| | GRNBoost2 | $\leq 0.001$ |
| CRC T-cells Zhang et al. and Li et al. | PPCOR | NA |
| | GENIE3 | $\leq 0.001$ |
| | GRNBoost2 | $\leq 0.001$ |
| HSC Hay et al. and Setty et al. | PPCOR | NA |
| | GENIE3 | $\leq 0.001$ |
| | GRNBoost2 | $\leq 0.001$ |
| CLP Hay et al. and Setty et al. | PPCOR | NA |
| | GENIE3 | $\leq 0.001$ |
| | GRNBoost2 | $\leq 0.001$ |
| Monocytes Hay et al. and Setty et al. | PPCOR | NA |
| | GENIE3 | $\leq 0.001$ |
| | GRNBoost2 | $\leq 0.001$ |
| Erythroblast Hay et al. And Setty et al. | PPCOR | NA |
| | GENIE3 | $\leq 0.001$ |
| | GRNBoost2 | $\leq 0.001$ |
| Dendritic Cell Hay et al. and Setty et al. | PPCOR | NA |
| | GENIE3 | $\leq 0.001$ |
| | GRNBoost2 | $\leq 0.001$ |

**Supplementary Table 3.** Number of links obtained for different subsamplings of the human retina dataset (Menon et al., 2019)

| Number of cells in subsampling | Algorithm | Number of links dataset 1 | Number of links dataset 2 |
| --- | --- | --- | --- |
| 10000 | PPCOR | 4963433 | 4966521 |
|  | GENIE3 | 2417821 | 2417821 |
|  | GRNBoost2 | 1959586 | 1971026 |
| 1000 | PPCOR | 645320 | 646830 |
|  | GENIE3 | 2417602 | 2417653 |
|  | GRNBoost2 | 1462312 | 1391567 |
| 100 | PPCOR | NA | NA |
|  | GENIE3 | 1987003 | 2157024 |
|  | GRNBoost2 | 666438 | 959804 |

## Notes

https://github.com/ComputationalSystemsBiology/scNET

### Competing Interest Statement

The authors have declared no competing interest.

## References

Aerts, S., Quan, X.-J., Claeys, A., Naval Sanchez, M., Tate, P., Yan, J., et al. (2010). Robust target gene discovery through transcriptome perturbations and genome-wide enhancer predictions in Drosophila uncovers a regulatory basis for sensory specification. PLoS biology 8, e1000435. doi:10.1371/journal.pbio.1000435.

Aibar, S., González-Blas, C. B., Moerman, T., Huynh-Thu, V. A., Imrichova, H., Hulselmans, G., et al. (2017). SCENIC: single-cell regulatory network inference and clustering. Nature Methods 14, 1083–1086. doi:10.1038/nmeth.4463.

Barabási, A.-L., Gulbahce, N., and Loscalzo, J. (2011). Network medicine: a network-based approach to human disease. Nat. Rev. Genet. 12, 56–68. doi:10.1038/nrg2918.

Barabási, A.-L., and Oltvai, Z. N. (2004). Network biology: understanding the cell’s functional organization. Nat. Rev. Genet. 5, 101–113. doi:10.1038/nrg1272.

Basso, K., Margolin, A. A., Stolovitzky, G., Klein, U., Dalla-Favera, R., and Califano, A. (2005). Reverse engineering of regulatory networks in human B cells. Nature Genetics 37, 382–390. doi:10.1038/ng1532.

Chan, T. E., Stumpf, M. P. H., and Babtie, A. C. (2017). Gene Regulatory Network Inference from Single-Cell Data Using Multivariate Information Measures. Cell Systems 5, 251–267.e3. doi:10.1016/j.cels.2017.08.014.

Chawla, K., Tripathi, S., Thommesen, L., Lægreid, A., and Kuiper, M. (2013). TFcheckpoint: a curated compendium of specific DNA-binding RNA polymerase II transcription factors. Bioinformatics 29, 2519–2520. doi:10.1093/bioinformatics/btt432.

Chen, S., and Mar, J. C. (2018). Evaluating methods of inferring gene regulatory networks highlights their lack of performance for single cell gene expression data. BMC bioinformatics 19, 232. doi:10.1186/s12859-018-2217-z.

Fiers, M. W. E. J., Minnoye, L., Aibar, S., Bravo González-Blas, C., Kalender Atak, Z., and Aerts, S. (2018). Mapping gene regulatory networks from single-cell omics data. Briefings in Functional Genomics 17, 246–254. doi:10.1093/bfgp/elx046.

Greenfield, A., Madar, A., Ostrer, H., and Bonneau, R. (2010). DREAM4: Combining genetic and dynamic information to identify biological networks and dynamical models. PloS One 5, e13397. doi:10.1371/journal.pone.0013397.

Hay, S. B., Ferchen, K., Chetal, K., Grimes, H. L., and Salomonis, N. (2018). The Human Cell Atlas bone marrow single-cell interactive web portal. Experimental Hematology 68, 51–61. doi:10.1016/j.exphem.2018.09.004.

Huynh-Thu, V. A., Irrthum, A., Wehenkel, L., and Geurts, P. (2010). Inferring regulatory networks from expression data using tree-based methods. PloS One 5. doi:10.1371/journal.pone.0012776.

Ideker, T., and Sharan, R. (2008). Protein networks in disease. Genome Research 18, 644–652. doi:10.1101/gr.071852.107.

Kim, S. (2015). ppcor: An R Package for a Fast Calculation to Semi-partial Correlation Coefficients. Communications for Statistical Applications and Methods 22, 665–674. doi:10.5351/CSAM.2015.22.6.665.

Li, H., Courtois, E. T., Sengupta, D., Tan, Y., Chen, K. H., Goh, J. J. L., et al. (2017). Reference component analysis of single-cell transcriptomes elucidates cellular heterogeneity in human colorectal tumors. Nat. Genet. 49, 708–718. doi:10.1038/ng.3818.

Lukowski, S. W., Lo, C. Y., Sharov, A. A., Nguyen, Q., Fang, L., Hung, S. S., et al. (2019). A single-cell transcriptome atlas of the adult human retina. EMBO J 38. doi:10.15252/embj.2018100811.

Margolin, A. A., Nemenman, I., Basso, K., Wiggins, C., Stolovitzky, G., Dalla Favera, R., et al. (2006). ARACNE: an algorithm for the reconstruction of gene regulatory networks in a mammalian cellular context. BMC bioinformatics 7 Suppl 1, S7. doi:10.1186/1471-2105-7-S1-S7.

Matsumoto, H., Kiryu, H., Furusawa, C., Ko, M. S. H., Ko, S. B. H., Gouda, N., et al. (2017). SCODE: an efficient regulatory network inference algorithm from single-cell RNA-Seq during differentiation. Bioinformatics 33, 2314–2321. doi:10.1093/bioinformatics/btx194.

Menon, M., Mohammadi, S., Davila-Velderrain, J., Goods, B. A., Cadwell, T. D., Xing, Y., et al. (2019). Single-cell transcriptomic atlas of the human retina identifies cell types associated with age-related macular degeneration. Nat Commun 10, 4902. doi:10.1038/s41467-019-12780-8.

Moerman, T., Aibar Santos, S., Bravo González-Blas, C., Simm, J., Moreau, Y., Aerts, J., et al. (2019). GRNBoost2 and Arboreto: efficient and scalable inference of gene regulatory networks. Bioinformatics (Oxford, England) 35, 2159–2161. doi:10.1093/bioinformatics/bty916.

Pratapa, A., Jalihal, A. P., Law, J. N., Bharadwaj, A., and Murali, T. M. (2020). Benchmarking algorithms for gene regulatory network inference from single-cell transcriptomic data. Nat. Methods 17, 147–154. doi:10.1038/s41592-019-0690-6.

Setty, M., Kiseliovas, V., Levine, J., Gayoso, A., Mazutis, L., and Pe’er, D. (2019). Characterization of cell fate probabilities in single-cell data with Palantir. Nat Biotechnol 37, 451–460. doi:10.1038/s41587-019-0068-4.

Silverman, E. K., Schmidt, H. H. H. W., Anastasiadou, E., Altucci, L., Angelini, M., Badimon, L., et al. (2020). Molecular networks in Network Medicine: Development and applications. Wiley Interdisciplinary Reviews. Systems Biology and Medicine, e1489. doi:10.1002/wsbm.1489.

Sonawane, A. R., Weiss, S. T., Glass, K., and Sharma, A. (2019). Network Medicine in the Age of Biomedical Big Data. Frontiers in Genetics 10, 294. doi:10.3389/fgene.2019.00294.

Tantardini, M., Ieva, F., Tajoli, L., and Piccardi, C. (2019). Comparing methods for comparing networks. Scientific Reports 9, 17557. doi:10.1038/s41598-019-53708-y.

Verny, L., Sella, N., Affeldt, S., Singh, P. P., and Isambert, H. (2017). Learning causal networks with latent variables from multivariate information in genomic data. PLoS Comput Biol 13, e1005662. doi:10.1371/journal.pcbi.1005662.

Zhang, Y., Zheng, L., Zhang, L., Hu, X., Ren, X., and Zhang, Z. (2019). Deep single-cell RNA sequencing data of individual T cells from treatment-naïve colorectal cancer patients. Sci Data 6, 131. doi:10.1038/s41597-019-0131-5.

